# Combination siRNA delivery as a therapeutic strategy for ADPKD

**DOI:** 10.64898/2026.07.20.739663

**Authors:** Joshua Giblin, Isabella Suzuki, Yi Huang, Karla Lambaren, Ronnie LaMastro, Rowan Simon, Jose Zarate-Diaz, Ali Osouli, Jessica Pham, Soobeen Laura Bae, Kenneth Hallows, Eun Ji Chung

**Affiliations:** Alfred E. Mann Department of Biomedical Engineering, University of Southern California, Los Angeles, California 90089, USA; Department of Medicine, Division of Nephrology, University of Vermont Health, Burlington, Vermont, 05405, USA; Silver Spur Therapeutics, Inc, Palos Verdes Peninsula, CA, 90274, USA; Department of Medicine, Division of Nephrology and Hypertension, Keck School of Medicine, University of Southern California, Los Angeles, California 90089, USA; Department of Chemical Engineering and Materials Science, University of Southern California, Los Angeles, California 90089, USA; Department of Surgery, Division of Vascular Surgery and Endovascular Therapy, Keck School of Medicine, University of Southern California, Los Angeles, California 90089, USA; Alfred E. Mann School of Pharmacy and Pharmaceutical Sciences, University of Southern California, Los Angeles, California 90089, USA

**Keywords:** Micelles, Gene Therapy, siRNA, ADPKD, Nanomedicine, Drug Delivery, Chronic Kidney Disease

## Abstract

Autosomal dominant polycystic kidney disease (ADPKD) is the most common genetic kidney disease worldwide, characterized by progressive cyst growth and inflammation, yet effective targeted therapies remain limited. Here we show that TMEM16A and MCP-1, key mediators of cyst-lining epithelial expansion and inflammatory macrophage recruitment respectively, are consistently upregulated in cyst-lining collecting duct (CD) epithelia across murine, porcine, and human ADPKD models. In human ADPKD patient cells, although individual silencing of *TMEM16A* or *MCP-1* transcripts reduced cyst growth, combined silencing produced superior therapeutic efficacy, establishing the rationale for evaluating dual-target delivery. To achieve dual gene silencing in the kidneys, we delivered *Tmem16a* and *Mcp*-1 siRNA using peptide amphiphile micelles (PAMs), an ultrasmall nanoparticle platform that enables efficient renal targeting. To redirect siRNA-loaded PAMs to CD epithelia, we functionalized their surface with a CD-targeting peptide (CDM), which enabled preferential accumulation in cyst-lining CD epithelia. In an inducible *Pkd1*-deficient mouse model, co-delivery of CDMs loaded with *Tmem16a* and *Mcp-1* siRNA reduced kidney enlargement, cystic burden, tubular injury, and macrophage infiltration, with efficacy exceeding non-targeted siRNA delivery at equivalent doses. CDM demonstrated enhanced uptake in primary human ADPKD patient-derived CD cells and dual gene silencing reduced target gene expression and cyst expansion, establishing translational relevance. These findings establish CD peptide–functionalized micelles as a route to cell-type-selective RNAi in the kidney, delivering siRNA to cyst-lining CD cells. Furthermore, because both targets, *TMEM16A* and *MCP-1*, are transcribed within CD cells, our siRNA-loaded CD-targeting micelles silence two drivers of cyst expansion, and their simultaneous suppression represents an effective therapeutic strategy for ADPKD.

## Introduction

Autosomal dominant polycystic kidney disease (ADPKD) affects 1 in every 400-1000 individuals, and is the most prevalent genetic kidney disease globally^1^. ADPKD is primarily caused by mutations in the *PKD1* (80% of cases) or *PKD2* (15% of cases) genes that encode the polycystin-1 (PC1) and polycystin-2 (PC2) proteins, respectively^2^. The loss of functional PC1 or PC2 in renal tubular epithelial cells results in the formation of fluid-filled cysts, which ultimately leads in patients reaching end-stage renal disease (ESRD) by 60 years of age^3^. Despite its prevalence, therapeutic options remain limited. The only FDA-approved treatment, tolvaptan, slows cyst growth by antagonizing vasopressin V2 receptor (V2R)–mediated cyclic adenosine monophosphate (cAMP) signaling in collecting duct (CD) epithelia, thereby reducing cell proliferation and fluid secretion that drive cyst expansion^4^. Because tolvaptan targets a single signaling axis within a multifactorial disease, its clinical benefit is modest, only delaying the onset of end-stage renal disease (ESRD) by an average of 3.1 years across chronic kidney disease (CKD) stages^4^. Furthermore, tolvaptan use is frequently associated with many undesirable side effects, including polyuria, excessive thirst, and hepatotoxicity, resulting in 25% of patients discontinuing its use^5,6,7,8,9,10,11^. These limitations highlight the need for therapies that simultaneously silence multiple disease-driving pathways.

Cyst expansion in ADPKD is driven by multiple, interrelated processes downstream of polycystin loss, including heightened proliferation, dysregulated fluid secretion and a pro-inflammatory microenvironment^12–14^. While inhibition of individual pathways reduces cyst burden in preclinical models, efficacy remains partial, because these processes operate in parallel rather than in series, suppressing one leaves the others intact^15,16^. This is analogous to that of cancer, where combination therapies targeting complementary mechanisms, such as proliferation and immune signaling, have demonstrated superior efficacy compared to single-agent approaches. To suppress multiple pathways driving cyst enlargement, we selected two genes, each controlling a distinct process of cyst growth, upregulated in cyst-lining epithelium, and individually validated as targets in preclinical ADPKD.

The first is TMEM16A or anoctamin 1 (ANO1), a calcium activated chloride channel upregulated in cyst-lining cells, which drives cyst enlargement through elevating chloride-mediated fluid secretion into the cyst lumen, promoting proliferation of cyst-lining cells, and promoting cilia dysfunction^12,17^. We selected TMEM16A rather than CFTR, another cAMP-activated chloride channel, because CFTR inhibition alone is insufficient to arrest cyst growth, and because TMEM16A acts upstream of CFTR^15,18^. Both channels are upregulated and colocalized in cyst-lining epithelia of *Pkd1*-deficient kidneys, but the cystic upregulation of CFTR is entirely reversed by deletion of *Tmem16a*, establishing that TMEM16A is required for proper expression and function of CFTR^15^. Moreover, deletion of *Tmem16a* restores cyst-lining cell proliferation to near-normal levels^15^. Therefore, silencing a single transcript, in *TMEM16A*, suppresses two arms of cyst enlargement, namely chloride secretion and cyst-lining proliferation.

For inhibiting a second major driver of PKD, inflammation, we selected monocyte chemoattractant protein 1 (MCP-1), a chemokine secreted by cyst-lining epithelia that recruits pro-inflammatory macrophages into the pericystic microenvironment through its receptor, C-C chemokine receptor 2 (CCR2), where they amplify cyst-lining cell proliferation through paracrine signaling^19^. The MCP-1/CCR2 axis has been validated at three independent points in preclinical PKD research. Notably, macrophage depletion with clodronate liposomes reduced cyst burden^20^, pharmacological antagonism of CCR2 attenuated cyst growth, and genetic deletion of *Mcp-1* reduced macrophage-dependent cyst expansion^16^. Furthermore, MCP-1 is elevated in the urine of ADPKD patients, where it correlates with disease severity^19^. Therefore, silencing MCP-1 can suppress both macrophage infiltration and the resulting inflammation in cystic kidneys.

Genetic disruption of either TMEM16A or MCP-1 alone reduces cyst burden in preclinical models of ADPKD, yet the kidneys remain significantly more cystic than healthy controls^15,16^. Residual disease with single-target disruption indicates that neither process alone accounts for the full cystic phenotype. As in the combination regimens that transformed cancer therapy, where combining agents against non-overlapping pathways can outperform single-agent therapy, we hypothesized that co-silencing TMEM16A, which controls fluid secretion and cyst-lining epithelial proliferation, and MCP-1, which controls macrophage recruitment and inflammation, would attenuate cyst progression more effectively than inhibiting either process alone^16,17^.

To silence TMEM16A and MCP-1, we chose to employ RNA interference (RNAi) using small interfering RNAs (siRNAs), a clinically validated and highly specific strategy to silence disease-driving genes. However, the clinical translation of siRNA therapeutics for kidney diseases remains limited by its biodistribution. Current clinically utilized siRNA systems, such as lipid nanoparticles (LNPs) and GalNAc-conjugated siRNAs are largely restricted to hepatic delivery and lack targeting to specific renal cell populations^21,22^. Thus, the development of kidney-targeted siRNA delivery platforms represents a critical unmet need, not only for ADPKD but for a broad range of renal diseases.

To enable kidney-specific delivery, we engineered ultrasmall nanoparticles decorated with a collecting duct targeting peptide to deliver siRNAs to the collecting duct (CD) epithelia. In addition to being the predominant cell type in which cysts arise, CD-derived cysts are considered the most disease-relevant in ADPKD, as multiple nephrons drain into each collecting duct, meaning that cyst formation at this convergence point obstructs flow from entire nephron networks, disproportionately impairing renal function relative to cysts arising elsewhere in the nephron^23^. Our prior work has shown that peptide amphiphile micelles (PAMs) composed of DSPE-PEG(2000) monomers demonstrate enhanced renal targeting abilities due to their small size (< 15 nm), enabling them to pass through the glomerular filtration barrier (GFB) and be taken up by renal tubular cells^24–26^. In addition to their passive renal targeting, the PEGylated lipid monomers can be functionalized with targeting peptides to promote tubule-specific cell uptake^24^. Here, we first verify that TMEM16A and MCP-1 are upregulated in cystic kidneys in murine, porcine, and human PKD samples. Moreover, in primary human ADPKD cells, combined knockdown of TMEM16A and MCP-1 produced greater suppression of cyst growth than either target alone, demonstrating complementary and cooperative contributions to disease progression. Building on this, we engineered CD-targeting micelles (CDM) functionalized with a collecting duct–targeting peptide, achieving enhanced uptake in CD cells in both cystic and healthy kidneys. In an inducible *Pkd1*-deficient mouse model, loading CDM with siRNA targeting *Tmem16a* and *Mcp-1* (CDM–siTM) reduced kidney enlargement, cystic burden, macrophage infiltration, and markers of renal injury compared to non-targeting controls. Critically, CDM–siTM reduced *TMEM16A* and *MCP-1* mRNA expression and decreased cyst expansion in primary human ADPKD patient cells, supporting clinical translation.

## Results

### TMEM16A and MCP-1 are upregulated in cyst-lining CD epithelia across preclinical models of ADPKD and their combined silencing suppresses cyst growth

To establish the therapeutic relevance of TMEM16A and MCP-1 as targets for ADPKD intervention, we first characterized their expression in cystic kidney tissue. To examine TMEM16A and MCP-1 expression in a mouse model of ADPKD, we used a well characterized, rapidly progressing inducible *Pkd1*^fl/fl^; Pax8rtTA;Tet-O-Cre mouse model, where mice are treated with doxycycline at postnatal (P) day 10 to induce *Pkd1* knockdown (**Fig. 1A**)^27^. On P21, disease progression was assessed, and induced mice (*Pkd1^-/-^*) exhibited a mean 6.1-fold higher kidney-weight-to-body weight (KW/BW) ratio compared to uninduced littermates (healthy, *Pkd1^+/+^*) (**Fig. 1A, B**). RT-qPCR of whole kidney lysate revealed an 8.8-fold increase in *Tmem16a* mRNA expression in *Pkd1^-/-^* mice compared to *Pkd1^+/+^* mice (**Fig. 1C**). Immunofluorescence co-staining with *Dolichos biflorus* agglutinin (DBA) revealed markedly elevated TMEM16A expression in DBA-positive cyst-lining CD cells of *Pkd1^-/-^*kidneys compared to healthy controls (**Fig. 1D**). To quantify this at the single-cell level, flow cytometric analysis of dissociated kidneys demonstrated that the proportion of TMEM16A-positive cells within the DBA-positive CD population increased from 13.1% in healthy kidneys to 47.3% in *Pkd1*^-/-^ kidneys, representing a 3.6-fold enrichment in TMEM16A-positive CD cyst-lining epithelia (**Fig. 1E, F**). We also examined *Mcp-1* mRNA and found that expression was 12.7-fold higher in *Pkd1^-/-^* kidneys compared to healthy kidneys (**Fig. 1G**). In addition, MCP-1 protein expression was markedly elevated in DBA-positive cyst-lining cells of *Pkd1*-deficient kidneys relative to healthy controls (**Fig. 1G**).

**Figure 1.**
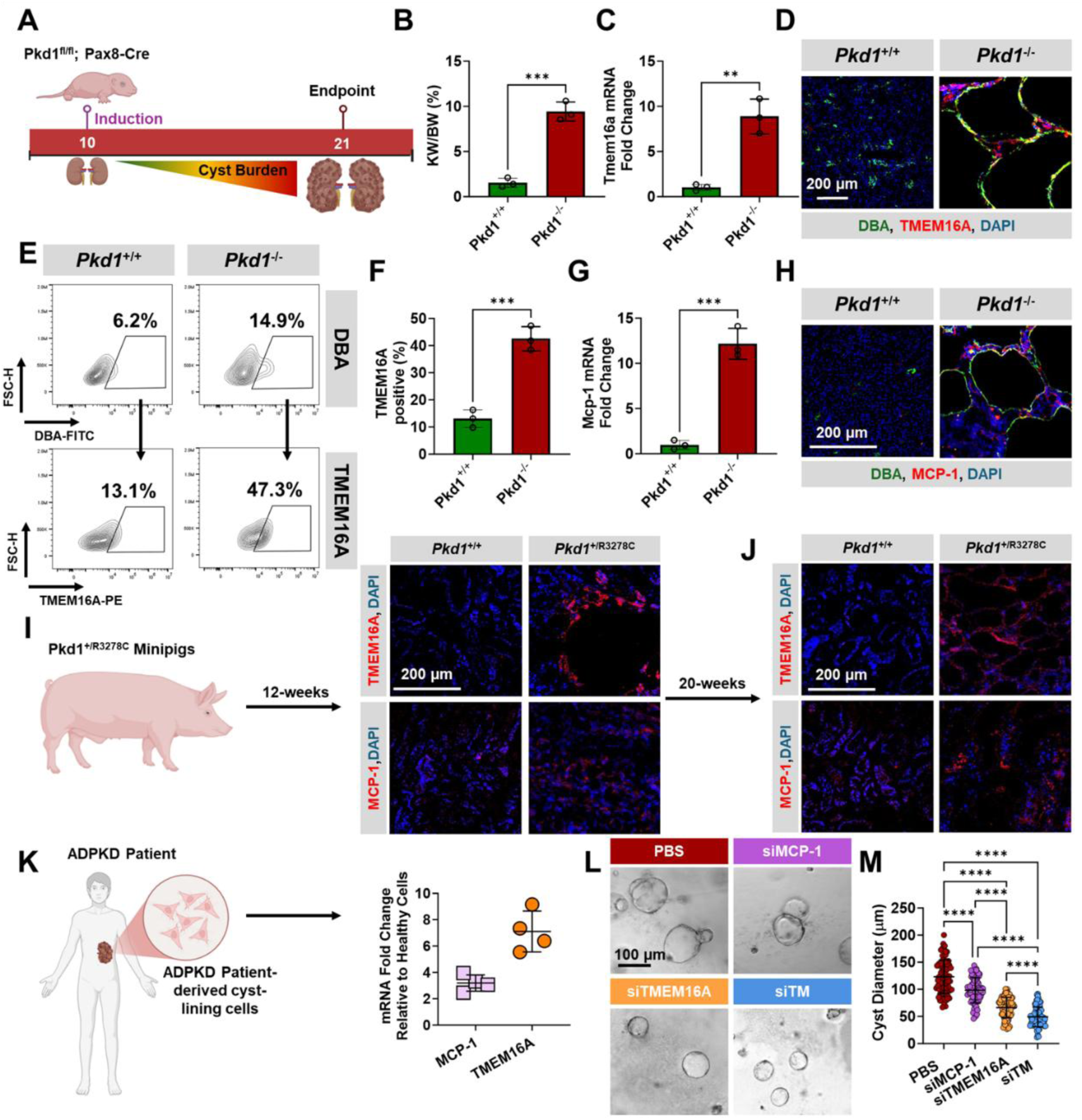
TMEM16A and MCP-1 are upregulated in cyst-lining CD epithelia across preclinical models of ADPKD and their combined silencing suppresses cyst growth. **(A)** Schematic of the inducible *Pkd1*^fl/fl^;Pax8-Cre mouse model. Mice were administered doxycycline at postnatal day 10 (P10) to induce *Pkd1* excision and were assessed at P21. **(B)** Kidney weight-to-body weight (KW/BW) ratios in *Pkd1*^+/+^ and *Pkd1*^-/-^ mice at endpoint (n = 3). **(C)** RT-qPCR quantification of *Tmem16a* mRNA expression in whole kidney lysates from both groups. **(D)** Representative immunofluorescence images of kidney sections co-stained for DBA (green), TMEM16A (red), and DAPI (blue) in both groups. Scale bar = 200 µm. **(E)** Representative flow cytometry plots showing gating strategy for DBA-positive CD cells and TMEM16A-positive cells within the DBA-positive population in *Pkd1*^+/+^ and *Pkd1*^-/-^ kidneys. **(F)** Quantification of the proportion of TMEM16A-positive cells within the DBA-positive CD population. **(G)** RT-qPCR quantification of *Mcp-1* mRNA expression in whole kidney lysates from both groups. **(H)** Representative immunofluorescence images of kidney sections co-stained for DBA (green), MCP-1 (red), and DAPI (blue). Scale bar = 200 µm. **(I)** Schematic of the *Pkd1*^+/R3278C^ minipig model with representative immunofluorescence images showing TMEM16A and MCP-1 expression in *Pkd1*^+/+^ and *Pkd1*^+/R3278C^ kidneys at 12 weeks. Scale bar = 200 µm. **(J)** Representative immunofluorescence images showing TMEM16A and MCP-1 expression in *Pkd1*^+/+^ and *Pkd1*^+/R3278C^ minipig kidneys at 20 weeks, demonstrating progressive upregulation with disease advancement. Scale bar = 200 µm. **(K)** Schematic of primary human ADPKD patient-derived cyst-lining cell isolation from nephrectomy specimens and RT-qPCR quantification of *MCP-1* and *TMEM16A* mRNA expression relative to healthy human renal cells (n = 4 patients). **(L)** Representative brightfield images of 3D cysts derived from human ADPKD patient cells treated with PBS, siMCP-1, siTMEM16A, or combined siRNA (siTM) beginning on day 3. Scale bar = 100 µm. **(M)** Quantification of cyst diameter at day 10 across treatment groups. Data are presented as mean ± SD. Statistical analysis was calculated with a one-way ANOVA with Tukey’s post hoc test if comparing more than two groups or student’s t-test if comparing two groups. ** p < 0.01, *** p < 0.001, **** p < 0.0001.

To determine whether this upregulation was conserved across species, we examined TMEM16A and MCP-1 expression in *Pkd1*^+/R3278C^ minipigs at 12- and 20-weeks of age. Both targets were elevated in cystic kidney tissue relative to healthy controls at both timepoints (**Fig. 1I, J**). Consistent with these findings, primary human ADPKD patient-derived cyst-lining cells exhibited 7.6-fold and 3.2-fold increases in *TMEM16A* and *MCP-1* mRNA expression, respectively, compared to healthy renal epithelial cells (**Fig. 1K**). To determine whether knockdown of these targets suppresses cyst growth as monotherapies or in combination, patient-derived ADPKD cells were cultured in a 3D cystogenesis assay and treated with si*TMEM16A*, si*MCP-1*, or both (siTM) beginning on day 3. Cysts treated with siTMEM16A and siMCP-1 were 47% and 21% smaller than PBS-treated controls by day 10, respectively (**Fig. 1L, M**). The more modest effect of siMCP-1 alone is consistent with its primary role in recruiting inflammatory macrophages to the pericystic niche rather than directly suppressing cystic cell proliferation, an effect that is limited in a monoculture cyst model lacking immune cell populations. Nonetheless, combination treatment with siTM produced the greatest reduction in cyst diameter (61%), supporting the rationale that dual targeting of proliferative/secretory and inflammatory/microenvironmental pathways provides therapeutic benefit beyond either monotherapy alone (**Fig. 1L, M**). Together, these findings establish that TMEM16A and MCP-1 are consistently upregulated in cyst-lining CD epithelia across murine, porcine, and human models of ADPKD, and that their simultaneous knockdown provides greater therapeutic benefit than either target alone, motivating the development of a CD-targeted delivery platform capable of achieving dual gene silencing *in vivo*.

### Decorating micelles with a collecting duct targeting peptide for enhanced targeting

Efficient siRNA delivery to cyst-lining CD epithelia requires a carrier capable of overcoming renal clearance barriers while achieving cell-type specific uptake^28^. We have previously developed PAMs composed of DSPE–PEG monomers, which self-assemble into ultrasmall nanostructures (6–20 nm) capable of traversing the GFB, a size constraint that limits conventional nanoparticle platforms such as lipid nanoparticles (LNPs), and accumulating within tubules following functionalization with targeting peptides^24,25^. To confer CD cell specificity, we incorporated a peptide ligand previously reported to bind to CD cells onto the micelle surface (**Fig. 2A**)^29^. The resulting CD-targeting micelles (CDMs) exhibited a spherical morphology with a hydrodynamic diameter of 14.5 ± 1.6 nm, a polydispersity index below 0.2, and a zeta potential of 3.7 ± 0.4 mV, consistent with a well-defined, monodisperse nanoparticle population (**Fig. S1A-D**)^24,26^. We next evaluated whether this targeting translates to preferential renal accumulation *in vivo*. *Pkd1^fl/fl^; Pax8rtTA;Tet-O-Cre* mice were administered doxycycline at postnatal day 10 (P10) to induce *Pkd1* excision and cyst formation^27^. At P18, mice received Cy7-labeled CDM or non-targeting micelles (NTM) via intraperitoneal (I.P.) injection and were euthanized 24 h later (P19).

**Figure 2:**
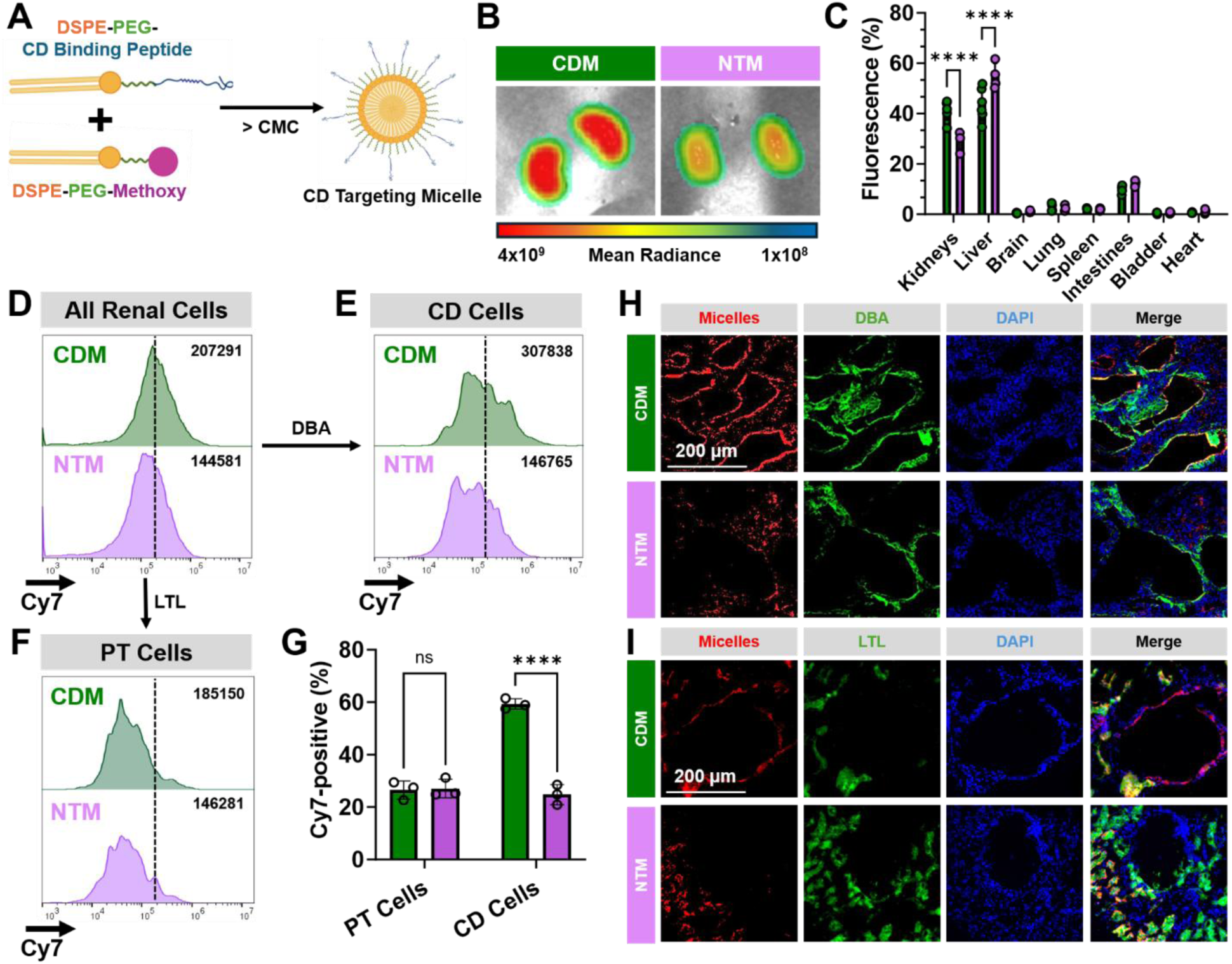
Collecting duct targeting micelles accumulate in collecting duct and cyst-lining epithelial cells *in vivo*. **(A)** Schematic of the collecting duct targeting micelles (CDM). **(B)** Representative *ex vivo* images showing micelle distribution in CDM-versus NTM-treated kidneys. **(C)** Quantification of Cy7 signal intensity across organs. Total photon flux was measured in the kidneys, liver, brain, lung, spleen, intestines, bladder and heart, and normalized by dividing the signal from each organ by the summed signal across all organs. **(D)** Flow cytometry histograms of fluorescence in all renal cells, comparing CDM and NTM. **(E)** Flow cytometry analysis of Dolichos biflorus agglutinin (DBA)-positive CD cells showing increased micelle association in CDM-treated mice relative to NTM. **(F)** Flow cytometry analysis of Lotus tetragonolobus lectin (LTL)-positive PT cells indicating comparable micelle accumulation between CDM and NTM treatments. **(G)** Quantification of micelle uptake across renal cell populations. **(H)** Representative confocal images of kidney sections from CDM- and NTM-treated mice stained for micelles (red), CD marker DBA (green), and nuclei (DAPI, blue), with merged images. Scale bar = 200 μm. **(I)** Representative confocal images of kidney sections from CDM- and NTM-treated mice stained for micelles (red), PT marker LTL (green), and nuclei (DAPI, blue), with merged images. Scale bar = 200 μm. Data are presented as mean ± SD (n ≥ 3). Statistical analysis was calculated with a student’s t-test if comparing two groups. **** p < 0.0001, ns = not significant.

*Ex vivo* fluorescence imaging of whole kidneys revealed significantly increased Cy7 signal in CDM-treated mice compared to NTM-treated controls with quantification confirming a 1.6-fold increase in kidney-associated fluorescence in the CDM group (**Fig. 2B**, **Fig. S2A**). Systemic biodistribution analysis showed predominant accumulation of both formulations in the liver. However, CDM-treated mice displayed enhanced kidney accumulation, exhibiting a 1.4-fold increase in kidney signal relative to NTM-treated mice (**Fig. 2C**). Consistently, the kidney-to-liver ratio was significantly elevated in the CDM group, indicating enhanced renal enrichment *in vivo* (**Fig. S2B**).

To resolve intrarenal distribution at the cellular level, flow cytometry was performed on dissociated kidney single-cell suspensions. Across all viable renal cells, CDM-treated kidneys displayed a 31% increase in Cy7 intensity compared to NTM-treated controls, consistent with *ex vivo* whole organ imaging (**Fig. 2D**). Cell type-specific analysis revealed that Cy7 intensity was 2.1-fold higher in DBA-positive CD cells of CDM-treated kidneys compared to NTM controls (**Fig. 2E**). In contrast, LTL-positive PT cells showed no significant difference in Cy7 intensity between treatment groups (**Fig. 2F**), demonstrating that enhanced CDM uptake is selective for CD over PT epithelia. Consistent with this, the fraction of Cy7-positive cells was unchanged in LTL-positive PT populations, whereas DBA-positive CD cells exhibited a 2.7-fold increase in Cy7-positive cells following CDM treatment, providing convergent evidence of CD-cell-specific accumulation by CDM (**Fig. 2G**).

To spatially confirm these findings, confocal microscopy was performed on kidney sections co-stained with cell-type specific markers. CDM signal co-localized extensively with DBA-positive cystic epithelia, whereas NTM showed minimal overlap (**Fig. 2H**). Although both CDM and NTM were detected in LTL-positive PT cells, CDM exhibited markedly enhanced accumulation within cyst-lining structures compared to NTM (**Fig. 2I**), consistent with the flow cytometric findings. Together, these data demonstrate that CDM achieves enhanced accumulation in cyst-lining CD epithelia *in vivo*, outperforming NTM at both the organ and cellular level.

### CD-targeted siRNA delivery attenuates disease progression in an ADPKD mouse model

Building on the demonstration that CDM achieves accumulation in CD cells *in vivo*, we next functionalized the platform to enable therapeutic siRNA delivery. siRNA was modified with a 5′-thiol on the sense strand and conjugated to DSPE–PEG(2000)–maleimide to generate a lipid–siRNA amphiphile, which was subsequently incorporated into CDM (CDM–siRNA). To determine siRNA loading capacity, a 21-mer siRNA amphiphile was incorporated into CDM at 1, 2, 5, and 10 mol%. Gel shift analysis showed that micelles formulated at 1 mol% retained siRNA, whereas higher loadings (2, 5, and 10 mol%) resulted in detectable RNA migration, indicating reduced encapsulation stability (**Fig. S3A**). Transmission electron microscopy confirmed that CDM–siRNA formulated with 1 mol% DSPE-PEG(2000)-siRNA maintained their size, spherical morphology, and low polydispersity, comparable to empty CDM (**Fig. S3B-E**).

Having demonstrated that TMEM16A and MCP-1 are upregulated in cyst-lining CD epithelia (**Fig. 1C-G**) and that their combined silencing suppresses cyst growth *in vitro* (**Fig. 1L, M**), we next evaluated the therapeutic efficacy of CDMs loaded with *Tmem16a* and *Mcp-1* siRNA (CDM–siTM) *in vivo*. *Pkd1^fl/fl^; Pax8rtTA;Tet-O-Cre* mice were administered doxycycline at P10 to induce *Pkd1* knockdown. Beginning at P15, mice received I.P. injections every other day until P21 of: CDM-siTM (1 mg/kg of each siRNA), NTM-siTM (1 mg/kg of each siRNA), free *Tmem16a* and *Mcp-1* siRNA (Free siTM), or PBS (**Fig. 3A**).

**Figure 3:**
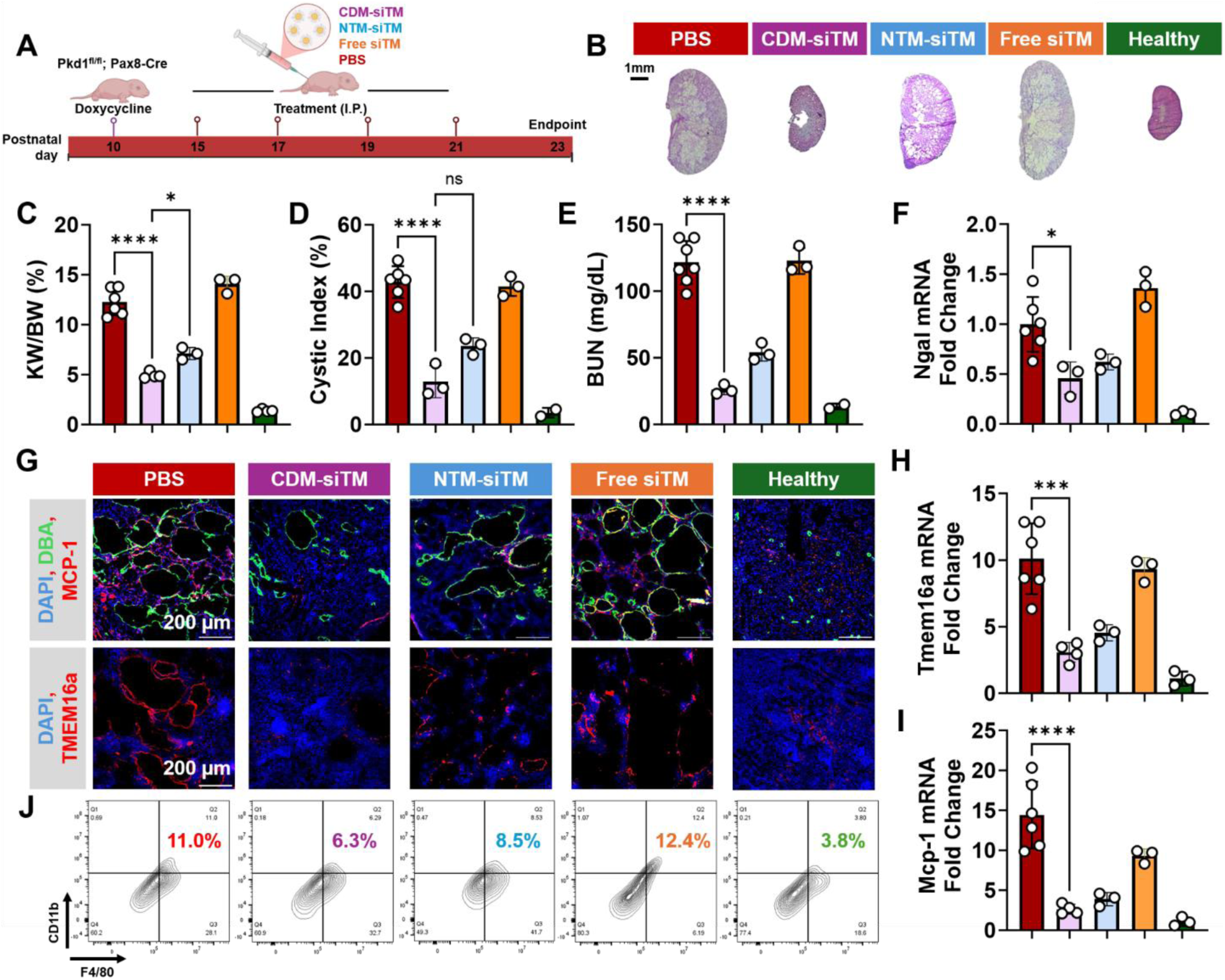
CDM-mediated dual siRNA delivery attenuates cyst growth and improves renal function in ADPKD. **(A)** Schematic of the *Pkd1*^fl/fl^; Pax8-Cre mouse model, including doxycycline induction at P10 and I.P. treatment schedule with CDM–siTM, NTM–siTM, free siTM or PBS, and endpoint analysis. **(B)** Representative kidney images at endpoint from each treatment group. Scale bar = 1 mm. **(C)** Kidney weight to body weight ratio (KW/BW) across treatment groups. **(D)** Quantification of cystic index across treatments. **(E)** Blood urea nitrogen (BUN) levels as a measure of renal function across groups. **(F)** Quantification of *Ngal* mRNA expression (fold change). **(G)** Representative immunofluorescence images of kidney sections from treated mice stained for MCP-1 (top row, red) and TMEM16A (bottom row, red), CD marker DBA (green), and nuclei (DAPI, blue). Scale bars = 200 µm. **(H)** RT-qPCR assessing *Tmem16a* mRNA and **(I)** *Mcp-1* mRNA across treatment groups. **(J)** Flow cytometry analysis from kidneys put into single cell suspension and stained with F4/80 and CD11b across treatments. Data are presented as mean ± SD (n ≥ 2). Statistical analysis was calculated with a one-way ANOVA with Tukey’s post hoc test. *** p < 0.001, **** p < 0.0001, ns = not significant.

At the study endpoint, gross morphology and histological analysis revealed attenuation of kidney enlargement in the CDM-siTM-treated cohort (**Fig. 3B**). CDM–siTM reduced the kidney weight-to-body weight (KW/BW) ratio by 60% relative to PBS-treated mice (**Fig. 3C**). Importantly, CDM–siTM reduced KW/BW by 29% compared to NTM–siTM-treated mice at the same dose, demonstrating that CD-specific targeting provides therapeutic benefit beyond passive micelle accumulation alone (**Fig. 3C**). In addition to reducing kidney enlargement, CDM–siTM reduced cystic index by 69% compared to PBS-treated controls and by 46% relative to NTM–siTM at the same dose, further supporting the importance of CD-targeted delivery for maximal therapeutic efficacy (**Fig. 3D**). Renal function assessed by blood urea nitrogen (BUN) was significantly improved in CDM–siTM-treated mice relative to PBS controls (**Fig. 3E**), indicating that structural improvements translated to functional benefit. Consistent with this, whole kidney *Ngal* expression^30,31^, a marker of tubular injury, was reduced by 51% in CDM–siTM-treated mice compared to PBS controls (**Fig. 3F**).

To define the mechanisms underlying the observed therapeutic effects, we examined target gene silencing and inflammatory signaling in kidneys from the treatment cohorts. Immunofluorescence analysis of kidney sections showed reduced MCP-1 protein levels in CDM–siTM-treated mice compared to PBS-treated controls, both in pericystic regions surrounding DBA-positive CD cells and throughout the kidney (**Fig. 3G**). In contrast, NTM–siTM-treated mice exhibited partial reduction with persistent MCP-1 protein expression in cyst-lining regions, consistent with the importance of CD-targeted delivery for maximal gene silencing in cyst-lining epithelia (**Fig. 3G**). RT–qPCR analysis confirmed an 82% reduction in *Mcp-1* mRNA in CDM–siTM-treated mice relative to PBS-treated controls; however, transcript levels remained elevated 2.6-fold compared to healthy kidneys (**Fig. 3H**). Similarly, TMEM16A protein expression was reduced in CDM–siTM-treated mice relative to PBS controls, with RT–qPCR revealing a 70% decrease in *Tmem16a* mRNA (**Fig. 3G, I**). Despite this significant reduction, *Tmem16a* expression remained 3.1-fold higher than in healthy tissue (**Fig. 3G, I**). Given the established role of MCP-1 in macrophage recruitment to the pericystic niche, we next assessed renal macrophage infiltration^16,32–35^. Flow cytometric analysis showed that PBS-treated mice exhibited a 2.9-fold increase in CD11b^+^F4/80^+^ macrophages compared to healthy controls (**Fig. 3J**). CDM–siTM treatment reduced macrophage infiltration by 43% relative to PBS-treated mice, whereas NTM–siTM produced a more modest reduction (23%), further supporting the role of CD-targeted MCP-1 silencing in attenuating inflammatory macrophage recruitment (**Fig. 3J**).

To evaluate systemic tolerability, body weight was monitored throughout the treatment period and remained stable across all groups (**Fig. S4A**). Additionally, CDM–siTM-treated mice showed no significant elevation in serum liver enzymes relative to PBS-treated controls or healthy mice at either dose (**Fig. S4B, C**). Histological examination of liver, lung, spleen, and heart revealed no evidence of tissue injury (**Fig. S4D**), indicating that repeated systemic administration of CDM–siTM at therapeutically effective doses is well tolerated. Together, these data demonstrate that CDM–siTM reduces TMEM16A and MCP-1 expression in cystic kidneys, attenuates macrophage infiltration and epithelial proliferation, and is well tolerated systemically, providing mechanistic support for the therapeutic effects observed at the organ and tissue level.

### CD-targeted siRNA delivery reduces inflammatory signaling, cellular proliferation, and cyst growth in ADPKD patient cells

To evaluate the translational relevance of CDM–siTM and determine whether the therapeutic effects observed in murine models extend to human disease, we assessed CDM targeting, functional gene silencing, and cystic phenotype suppression in primary human ADPKD patient-derived cells isolated from cysts. Flow cytometric analysis confirmed that patient-derived cells stained positive for DBA and negative for LTL, establishing their identity as CD epithelia rather than PT cells (**Fig. S5**)^36^. Consistent with this, CDM exhibited greater association with patient-derived ADPKD cells than NTM, confirming that peptide-mediated CD targeting is preserved in human disease-relevant cells (**Fig. 4A**).

**Figure 4:**
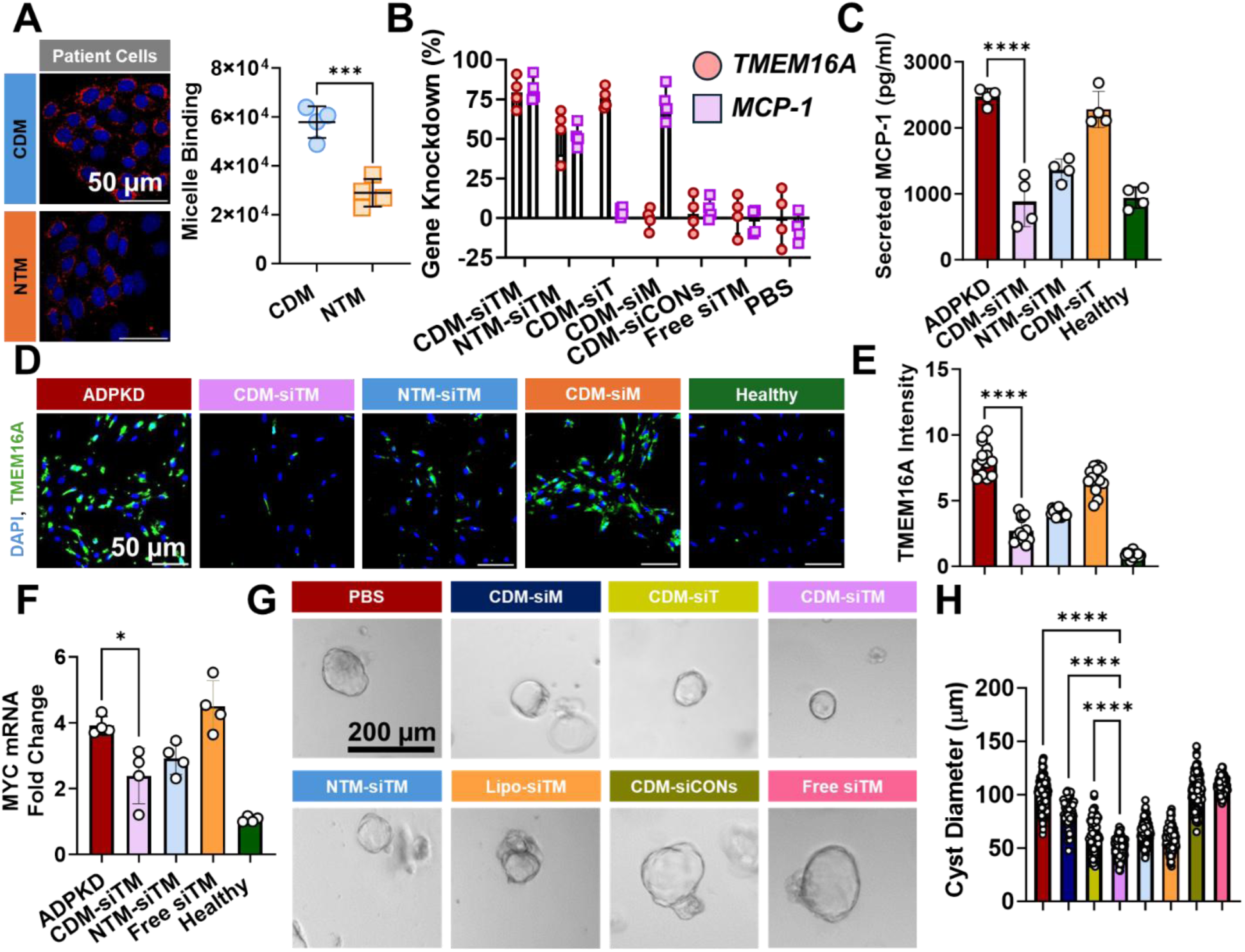
CDM-mediated dual gene silencing reduces inflammation, proliferation, and cyst growth in ADPKD patient cells. **(A)** Fluorescence images and quantification of Rhodamine B-labeled CDM or NTM (red) uptake in primary human ADPKD cells. **(B)** RT–qPCR analysis of *TMEM16A* and *MCP-1* mRNA expression in primary human ADPKD cells relative to healthy renal cells after treatment with CDM–siTM, NTM–siTM, CDM–siT, CDM–siM, CDM–siCONs, free siTM, or PBS. **(C)** MCP-1 secretion measured in culture supernatants following treatment of ADPKD cells with the indicated formulations. **(D)** Representative immunofluorescence images of TMEM16A (green) and nuclei (DAPI, blue) in ADPKD, CDM–siTM, NTM–siTM, CDM–siM, and healthy cells. Scale bar = 50 µm. **(E)** Quantification of TMEM16a fluorescence intensity. **(F)** RT–qPCR analysis of *MYC* mRNA expression across the indicated groups. **(G)** Representative brightfield images of 3D cysts treated with PBS, CDM-siM, CDM-siT, CDM–siTM, NTM–siTM, Lipo–siTM, CDM–siCONs, or free siTM. Scale bars = 200 µm. **(H)** Quantification of cyst diameter across treatment groups. Each point represents an individual cyst. Data are presented as mean ± SD, (n ≥ 4). Statistical analysis was calculated with a one-way ANOVA with Tukey’s post hoc test. *** p < 0.001, **** p < 0.0001, ns = not significant.

Treatment of ADPKD patient-derived cells with CDM loaded with human-specific siT, siM, or both (siTM), for 48 h resulted in significant knockdown of both transcripts as measured by RT–qPCR (**Fig. 4B**). CDM–siTM achieved greater suppression of *TMEM16A* and *MCP-1* mRNA than NTM–si*TM*, which showed 14% lower knockdown efficiency, confirming that CD-targeted delivery enhances gene silencing beyond passive micelle accumulation in human ADPKD cells (**Fig. 4B**). At the protein level, MCP-1 secretion into the culture medium was reduced to levels comparable to those observed in healthy renal cells following CDM–siTM treatment (**Fig. 4C**). Immunofluorescence analysis confirmed CDM–siTM produced the greatest reduction relative to untreated ADPKD cells (**Fig. 4D, E**). In addition, CDM–siTM reduced *MYC* mRNA expression in ADPKD cells, a downstream effector of cyst-associated proliferative signaling, although levels remained elevated relative to healthy controls, consistent with partial but not complete suppression of cystogenic transcriptional programs (**Fig. 4F**).

To determine whether dual gene silencing translates to functional inhibition of cyst growth in human disease, ADPKD patient-derived CD cells were cultured in a 3D cystogenesis assay and treated with CDM–siTM, NTM–siTM, CDM–siCONs, Lipo–siTM, free siTM, or PBS from day 3 onwards (**Fig. 4G**). By day 10, CDM–siTM-treated cysts were 49% smaller in diameter than PBS controls, representing the greatest reduction across all treatment groups (**Fig. 4H**). Critically, CDM–siTM outperformed CDM–siM and CDM–siT monotherapies by 33% and 19%, respectively, confirming that simultaneous suppression of both TMEM16A and MCP-1 produces additive therapeutic benefit that neither target alone achieves, consistent with the mechanistic non-redundancy of epithelial fluid secretion and inflammatory macrophage recruitment as parallel drivers of cyst expansion. CDM–siCONs and free siTM produced negligible effects on cyst diameter relative to PBS controls, establishing that therapeutic efficacy requires the combination of nanoparticle-mediated delivery and sequence-specific gene silencing, and excluding non-specific nanoparticle or siRNA effects as confounders (**Fig. 4G, H**).

Together, these findings demonstrate that CDM achieves preferential association with human ADPKD patient-derived cells, enables efficient dual gene silencing of *TMEM16A* and *MCP-1*, and suppresses cyst expansion in a human 3D PKD cyst model, establishing the translational potential of CD-targeted PAMs for therapeutic siRNA delivery in ADPKD.

## Discussion

ADPKD is the most common monogenic cause of kidney failure, affecting approximately 1 in 400–1,000 individuals worldwide and accounting for 7-15% of patients requiring renal replacement therapy^23,37^. Despite decades of mechanistic investigation, tolvaptan remains the only FDA-approved disease-modifying therapy for ADPKD. However, its use is associated with serious adverse effects including hepatotoxicity, and it is limited to patients at risk of rapid disease progression^11^. Critically, tolvaptan targets a single pathway, vasopressin 2-driven cAMP signaling, and slows but does not halt disease progression, leaving most of the pathological landscape unaddressed. Despite active investigation of agents targeting mTOR signaling^38^, AMPK activation^39^, tyrosine kinase pathways^40^, and metabolic reprogramming^41^, none has demonstrated sufficient efficacy to achieve regulatory approval. Further, no current strategy has been granted FDA-approval, underscoring that slowing ADPKD progression through single-pathway inhibition alone may be insufficient, and that a fundamentally different therapeutic framework is required.

This therapeutic logic mirrors the combination therapy paradigm that transformed oncology, where the recognition that complex diseases resist single-pathway inhibition drove the development of regimens targeting complementary mechanisms simultaneously, a strategy that has repeatedly produced better outcomes that either agent achieves alone. ADPKD presents an analogous therapeutic challenge, where cyst growth is sustained by at least two mechanistically distinct processes^33,35,42,43^. TMEM16A (ANO1) is a calcium-activated chloride channel expressed on the apical surface of cyst-lining epithelia. TMEM16A drives luminal fluid secretion and epithelial proliferation and has been previously validated as a major contributor to cyst expansion in mouse models and human ADPKD^15,44^. In parallel, MCP-1 coordinates macrophage recruitment to the cyst microenvironment, establishing a pro-inflammatory niche that amplifies cyst growth through paracrine signaling independent of epithelial chloride transport^16,35^. Because these pathways operate in parallel rather than in series, targeting downstream pathways beyond a single mechanism may provide additive or synergistic therapeutic value, and neither monotherapy alone is sufficient to fully arrest disease progression. This is directly confirmed by our data, where CDM–siTM produced a 33% and 19% greater reduction in cyst diameter compared to CDM–siM and CDM–siT monotherapies, respectively, an additive benefit that validates the combination hypothesis.

Translating dual knockdown of *TMEM16A* and *MCP1* using siRNA requires overcoming a bottleneck in renal nucleic acid therapeutics, delivery. Despite six FDA-approved siRNA therapeutics, patisiran, givosiran, lumasiran, inclisiran, vutrisiran, and nedosiran, all target the hepatocytes of the liver^45^. Patisiran relies on LNP-mediated hepatic delivery via apolipoprotein E-mediated uptake, while the five subsequent approvals employ GalNAc conjugates that exploit the asialoglycoprotein receptor, which is highly expressed on hepatocytes^46^. The kidney presents a fundamentally different anatomical challenge. The glomerular filtration barrier acts as a size-selective gate with molecules below approximately ∼6 nm undergoing filtration and rapid urinary clearance, while those above approximately ∼15 nm are excluded from the tubular compartment entirely^47^. This narrow window excludes conventional nanoparticle platforms, including the lipid nanoparticles (LNPs) used in patisiran, which typically range from 60–100 nm, from accessing the renal filtration system. To date, no siRNA therapeutic has yet achieved clinical approval for any kidney disease, and no platform has demonstrated cell-type-specific delivery to CD epithelia, the primary site of cystogenesis in ADPKD.

We addressed this gap by engineering peptide amphiphile micelles (PAMs) that exploit the unique anatomy of the renal filtration system. By formulating DSPE-PEG monomers into ultrasmall nanostructures of 6–15 nm, a size regime specifically designed to traverse the glomerular filtration barrier and access the tubular lumen, and functionalizing the micelle surface with a CD-targeting peptide, we achieved delivery of siRNA to CD epithelia.

CDM-mediated dual silencing of TMEM16A and MCP-1 produced therapeutic outcomes that neither monotherapy achieved alone, suppressing cyst expansion in 3D PKD models and attenuating cyst burden and immune cell infiltration in *Pkd1*-deficient mice. The superiority of combination over monotherapy is consistent with the mechanistic non-redundancy of these two targets. Only by simultaneously engaging both arms does the platform produce the degree of therapeutic benefit observed in our studies, a finding that directly validates the combination therapy hypothesis and establishes a precedent for multi-target siRNA delivery in chronic kidney disease.

The demonstration of preferential CDM uptake in primary human ADPKD patient-derived CD cells and suppression of cyst expansion in human 3D cyst models strengthens the translational relevance of these findings, given the well-documented challenges of extrapolating murine PKD data to human disease. The marked upregulation of TMEM16A and MCP-1 in patient-derived cells relative to healthy controls, combined with the therapeutic response to CDM–siTM, confirms that both targets are clinically accessible and disease-relevant in human ADPKD.

Despite our promising results, several limitations warrant consideration. First, while CDM delivery was enriched in CD cells, uptake was also observed in PT and other intrarenal populations. Notably, MCP-1 expression in non-CD cell types, including pericystic fibroblasts and infiltrating macrophages, represents a disease-relevant rather than purely off-target population, and broader intrarenal distribution may provide complementary benefit. Second, the *in vivo* durability of gene knockdown under the dosing regimen employed remains to be directly characterized. While the full duration of CDM-siTM-mediated *Tmem16a* and *Mcp-1* mRNA silencing will be fully explored in future studies, previous studies by our group that utilized a similar PAM platform to deliver microRNA, have demonstrated sustained functional and therapeutic effects for up to two months following a single dose *in vivo*, suggesting that micelle-mediated nucleic acid delivery to kidney cells also has potential to support silencing durations within this range^48,49^. This duration aligns with clinically approved siRNAs, such as patisiran, which demonstrates knockdown of its target gene, transthyretin (*TTR)* for up to 2 months in non-human primates (NHP)^50^. Strategic incorporation of chemical modifications into the siRNA strands, including 2′-Ome and 2’-F substitutions, phosphorothioate linkages, and 5′-(E)-vinyl phosphonate modifications, may further enhance siRNA stability and extend the duration of target gene silencing once in the cytosol. Third, studies were conducted in a complete *Pkd1* inactivation model, which may not fully recapitulate the partial loss-of-function mutations characteristic of most ADPKD patients^30,51^. Thus, validation in hypomorphic models such as Pkd1^RC/RC^ will be important^52^. Finally, therapeutic initiation after established cyst burden, more representative of the clinical scenario, remains to be evaluated.

Together, these findings establish CD-targeted PAMs as the first platform to achieve mechanistically defined, cell-type-specific siRNA delivery to cyst-lining epithelia and demonstrate that simultaneous suppression of epithelial fluid secretion and inflammatory macrophage recruitment produces additive therapeutic benefit exceeding either approach alone. By resolving the kidney delivery problem that has limited siRNA therapeutics to hepatic targets, and by validating a dual-pathway silencing strategy grounded in the combination therapy logic that transformed oncology, this work provides both a platform and a conceptual framework for the next generation of precision RNA therapeutics in ADPKD and broader renal disease.

## Methods

### Materials

DSPE-PEG(2000)-amine, DSPE-PEG(2000)-methoxy, and DSPE-PEG(2000)-maleimide were purchased from Avanti Polar Lipids (Alabaster, AL, USA). Amino acids were purchased from Gyros Protein Technologies (Uppsala, Sweden). Cy7 mono-N-hydroxy succinimide (NHS) ester was purchased from Lumiprobe (Hunt Valley, MD, USA). *TMEM16A*, MCP1, and control siRNA duplexes were purchased from Millipore Sigma (Burlington, MA, USA). M1 CCD cells and primary renal proximal tubular epithelial cells (RPTECs) were purchased from American Type Culture Collection (ATCC, Manassas, VA, USA). Primary ADPKD patient cells were provided by the PKD research resource consortium (PKD RRC, University of Maryland, Baltimore). Cell culture reagents were purchased from Sigma Aldrich and Gibco (Waltham, MA, USA).

### Synthesis of Peptide Amphiphiles

The CD targeting peptide CKDSPKSSKSIRFIPVST was synthesized on an automated peptide synthesizer (PS3, Gyros Protein Technologies, Tucson, AZ) using standard Fmoc-mediated, solid phase peptide synthesis methods. To cleave the peptide from the resin, a 94:2.5:2.5:1 vol% of trifluoroacetic acid: 1,2-ethanedithiol: water: triisopropylsilane was added to the beads and reacted for 4h. Peptides were dried with nitrogen, precipitated, and washed twice with ice-cold diethyl ether. The crude peptide was dissolved and lyophilized. Lyophilized peptide was dissolved in milliQ water and purified on a reverse-phase high performance liquid chromatography system (HPLC, Prominence, Shimadzu, Columbia, MD) using a Luna C8 column (250 × 10 mm ID, 5 μm, Phenomenex, Torrance, CA) and a gradient mobile phase consisting of (A) 0.1% (v/v) formic acid in water and (B) 0.1% (v/v) formic acid in acetonitrile. The m/z was characterized by MALDI-TOF/TOF (Bruker, Billerica, MA, USA). The expected m/z peak is 2022 g/mol. The purified peptides were lyophilized and conjugated to DSPE-PEG2000-maleimide in pH 7.4 buffered milliQ water for 72h. The solution was then purified using a C4 Jupiter column (250 × 30 mm ID, 5 μm, Phenomenex, Torrance, CA) as described above. The expected m/z peak for the peptide amphiphiles are [M+H]+ = 4963.

### Synthesis of RNA Amphiphiles

*TMEM16a* siRNA and MCP1 siRNA were modified with a 5’ thiol on the sense strand and mixed with DSPE-PEG2000-maleimide in pH 7.2 buffered PBS for 24h to form a thioether bond.

*TMEM16a* siRNA Mouse: 5’ – UUUAGACAAAAACCAAUAGAU

*TMEM16a* siRNA Human: 5’ – GAAGCAACACCTATTCGACCTG

MCP1 siRNA Mouse: 5’ – CCGUAAAUCUGAAGCUAAUdTdT

MCP1 siRNA Human: 5’ – CTCGCGAGCTATAGAAGAA

### Micelle Synthesis

Peptide amphiphile micelles were assembled through thin-film self-assembly methods as previously described. Briefly, CD targeting peptide amphiphiles (CD1-4) were dissolved in methanol, the solvent completely evaporated under nitrogen, and the resulting film was left overnight under vacuum. Subsequently, the film was hydrated in PBS at 1/10^th^ the working volume for *in vitro* studies, milliQ water for characterization studies, or PBS for *in vivo* studies. The solution was vortexed for 10s, sonicated for 1h, and incubated at 60°C for 30m. All micelles were assembled at least 4h prior to use to allow for temperature equilibration. For fluorescence studies, DSPE-PEG2000-rhodamine B or DSPE-PEG2000-Cy7 was incorporated into micelles at 10 mol%. For CD3-siRNA or NT micelles, the peptide amphiphile was dissolved in methanol, the solvent was evaporated under nitrogen, and the film was left under vacuum overnight. DSPE-PEG2000-siRNA at 1mol% was diluted in sterile PBS/ethanol and added to the lipid film and incubated at 50°C for 30 min. siRNA micelles were then dialyzed against 1X PBS overnight.

### Dynamic Light Scattering and Zeta Potential

Hydrodynamic diameter, zeta potential, and polydispersity index were determined using a Malvern Zetasizer Ultra instrument (Malvern Panalytical, Malvern, UK) and measurements were carried out at room temperature in triplicates N=3.

### Transmission Electron Microscopy

Transmission Electron microscopy was used to observe the morphology and dispersity of CD targeting micelles as previously described^53^. Briefly, carbon grids were wetted with 5 µl of milliQ water for 5 min. 50 µM 0.2 µm PVDF filtered micelle solution was added onto the grids and allowed to rest for 5 min. 2 wt% uranyl acetate was added onto the grid for 3 min, twice. Samples were kept in the dark and imaged on a FEI Talos F200C microscope (Thermo Fisher).

### Biodistribution of PAMs in an ADPKD mouse model

*Pkd1*^fl/fl^;Pax8-rtTA;Tet-O-Cre mice were I.P. injected with 50 mg/kg of doxycycline on postnatal day 10. On day 18, the mice were I.P. injected with 100 µl of sterile PBS containing PAMs at 1 mM (10mol% DSPE-PEG(2000)-Cy7). 24 h later, the mice were euthanized by CO2 overdose, their organs removed and imaged using an IVIS. Following imaging, the organs were snap frozen in tissue OCT using liquid nitrogen. For flow cytometry, one kidney from each mouse was finely minced, placed in RPMI containing 1 mg/ml collagenase and agitated on a shaker for 60 min at 37°C. The tissue was passed through a 40 µm filter before being spun at 300 g for 5 min at 4°C. The pellet was suspended in flow cytometry buffer, incubated with antibodies for 30 min at 4°C, washed once more, and ran through a flow cytometer. The gating strategy is as follows: forward scatter (FSC) versus side scatter (SSC) plots were used to exclude debris from the analysis, and singlets were selected via FSC-H vs FSC-A plots. Then, Cy7-positive cells were identified by gating for Cy7 based on PBS-treated control mice^53,54^,.

### Immunohistochemistry

Kidneys were sectioned at 8 µm using a Leica cryostat, dipped in Tris buffered saline with 0.1% Tween-20 (TBST), fixed in 4% PFA, and permeabilized with TBS containing 0.1% Triton-X 100 for 30 minutes. The sections were then washed thrice with TBST, blocked with 5% normal goat serum and 2% BSA for 1h at RT, and washed thrice more. Following this, the sections were incubated with primary antibodies or pre-fluorescently labeled molecules overnight at 4°C. The following day, the sections were washed, incubated with secondary antibodies (1:1000) for 1h at RT, washed, counter stained with DAPI (1:1000) for 1h at RT, and washed once more. A glass coverslip was then mounted onto kidney sections using Vectamount mounting medium and imaged on an LSM 880 confocal microscope. All quantification of images is performed using ImageJ and data points are the average of > 6 kidney sections with > 3 images per section per mouse.

| Name | Dilution | Catalog Number |
| --- | --- | --- |
| Dolichos Biflorus Agglutinin (DBA), Fluorescein | 1:200 | Vector Laboratories (FL-1031-2) |
| Lotus Tetragonolobus Lectin (LTL), Fluorescein | 1:200 | Vector Laboratories (FL-1321-2) |
| F4/80 Monoclonal Antibody | 1:100 | ThermoFisher (MA5-16363) |
| Goat anti-Rabbit IgG (H+L) Cross-Adsorbed Secondary Antibody, Alexa Fluor™ 594 | 1:1000 | ThermoFisher (A-11012) |
| CD45 Monoclonal Antibody (30-F11), FITC | 1:100 | ThermoFisher (11-0451-82) |
| F4/80 Monoclonal Antibody (BM8), APC | 1:100 | ThermoFisher (17-4801-82) |
| CD31 (PECAM-1) Monoclonal Antibody (390), FITC | 1:200 | ThermoFisher (11-0311-82) |
| Wheat Germ Agglutinin (WGA)-Alexa Fluor 488 | 1:100 | ThermoFisher (W11261) |
| Anti mouse *TMEM16a* | 1:100 | Bioss - BS-3794R |
| Anti human *TMEM16a* | 1:100 | Sino Biological - 102411-T10 |
| Anti mouse MCP1 | 1:200 | Thermo - PA5-115555 |

### In vitro knockdown analysis of TMEM16a and MCP1 mRNA

IMCD *Pkd1* knockout (KO), IMCD WT, primary cyst-lining epithelial cells from ADPKD patients, or renal tubular epithelial cells from healthy kidneys were seeded at 150,000 cells/well in a 12-well plate, left to adhere overnight, and incubated with micelles containing siRNA. Following treatment, cells were lysed, RNA was isolated via mRNeasy kit from Qiagen (Hilden, DE), and cDNA was synthesized using the RT2 first strand kit according to manufacturer instruction (Qiagen). The expression of *TMEM16A*, MCP1, and GAPDH was evaluated using real-time PCR with primer assays (Qiagen) and the RT2 SYBR green qPCR mastermix (Qiagen) on a CFX384 Touch Real-Time PCR detection system (Bio-Rad Laboratories). Fold-change in mRNA expression was calculated using the delta-delta Ct method.

*Measuring* MCP-1 *secretion* MCP-1 protein expression was evaluated through an ELISA as previously described^34^. Briefly, cells were plated at a density of 50,000 cells/well in a 12-well plate and left to adhere overnight. Cells were washed and allowed to grow for another 48 h after which the cells were washed, and fresh medium was replenished. The cells grew for another 3 days, and medium was collected. Protein levels were normalized using a BCA protein assay. Each data point is the mean of triplicate measurements.

### Evaluating the therapeutic efficacy in an ADPKD mouse model

*Pkd1*^fl/fl^;Pax8-rtTA;Tet-O-Cre mice were I.P. injected with 50 mg/kg of doxycycline on postnatal day 10. Starting on day 15, mice were I.P. injected with treatments every other day until day 21. On day 23, the mice were euthanized by CO2 overdose and their organs collected.

### Histology and Immunohistochemistry

Fresh kidney sections were then stained with hematoxylin and eosin (H&E) or Sirius red, according to the manufacturer’s instructions. Images were captured using a Leica DMi8 microscope (Leica, Wetzlar, Germany). For immunohistochemistry, fresh sections were fixed with 10% formalin for 10 min, permeabilized with 0.1% triton-X100 for 30 min, and blocked with 2% BSA for 30 min. Following several washes, sections were incubated with primary antibodies against macrophages (F4/80, 1:100), *TMEM16a* (1:100), or MCP-1 (1:200) overnight at 4°C. The following day, samples were incubated with a fluorophore conjugated secondary antibody (1:1000) and DAPI (1:1000) for 1 h at room temperature. Images were captured using a Leica DMi8 microscope (Leica, Wetzlar, Germany).

### Serum Markers and Toxicity Assessment

Blood samples were collected at experimental endpoints. Serum levels of alanine aminotransferase (ALT; Sigma, Cat: MAK571), aspartate aminotransferase (AST; Sigma, Cat: MAK055), and blood urea nitrogen (BUN; Invitrogen, Cat: EIABUN) were measured according to the manufacturer’s instructions to evaluate liver and kidney function.

### Statistical Analysis

All experiments were performed with at least two biological replicates unless otherwise specified. Data are presented as mean ± standard deviation. Statistical analyses were performed using GraphPad Prism software. Comparisons between two groups were performed using unpaired two-tailed Student’s t-tests, while multiple group comparisons were conducted using one-way or two-way ANOVA with appropriate post hoc tests. Differences were considered statistically significant at p < 0.05.

## Acknowledgments

The authors would like to acknowledge the NIDDK O’Brien Kidney National Resource Center (U54DK137516-03), NIDDK ISAC Award Program (U24DK128851), Rosalie and Harold Rae Brown Charitable Foundation, USC Transformative Center for Nanomedicine, and Silver Spur Therapeutics, Inc. The authors would also like to thank the Center for Electron Microscopy and Microanalysis (CNI) and the Translation Imaging Center (TIC) at USC for assistance in TEM and confocal imaging. In addition, the authors would like to thank the PKD Research Resource Consortium (PKD-RRC) at the University of Maryland School of Medicine, Baltimore, MD for assistance in retrieving ADPKD patient cells. Finally, the authors would like to thank Exemplar Genetics (Sioux Center, IA, USA) for providing pig renal tissue.

---

**Figure S1:**
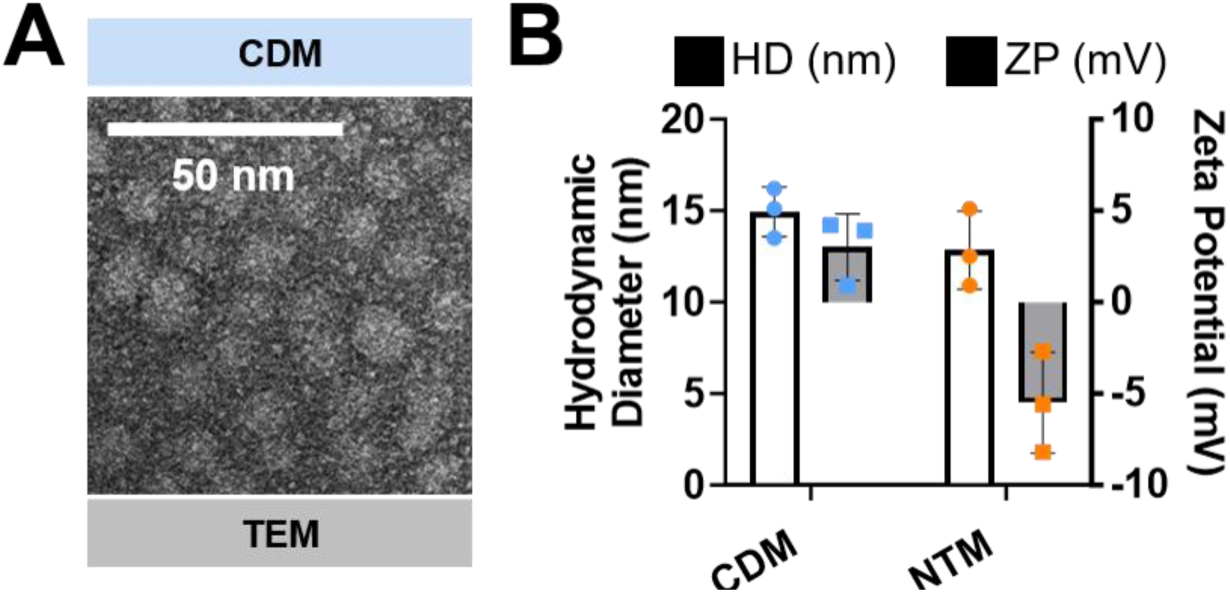
Characterization of collecting duct targeting micelles (CDMs). **(A)**Transmission electron microscopy (TEM) images demonstrate that CDMs exhibit a spherical morphology with a small, uniform size and a monodisperse distribution. Scale bar = 50 nm. **(B)** Dynamic light scattering (DLS) analysis of the hydrodynamic diameter (nm) and zeta potential (mV) of CDMs and non-targeting micelles (NTMs). Data are presented as mean ± SD (N = 3).

**Figure S2:**
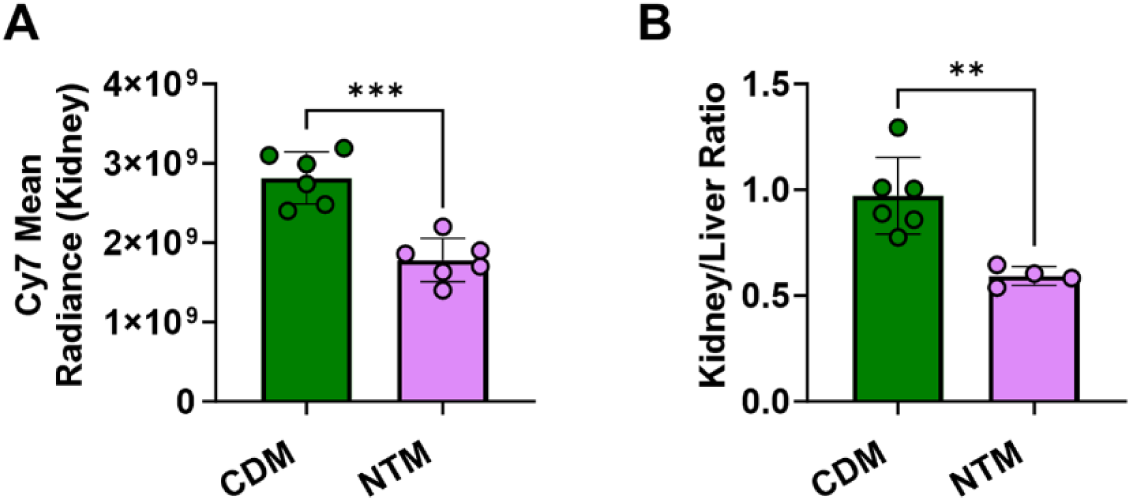
Quantification of renal accumulation and kidney-to-liver distribution of CD-targeted micelles. (A) Quantification of Cy7 fluorescence intensity in kidneys following administration of CDM or NTM. (B) Kidney-to-liver fluorescence ratio for CDM- and NTM-treated mice. (n ≥ 4). Data are presented as mean ± SD with individual data points representing biologically independent animals.

**Fig. S3:**
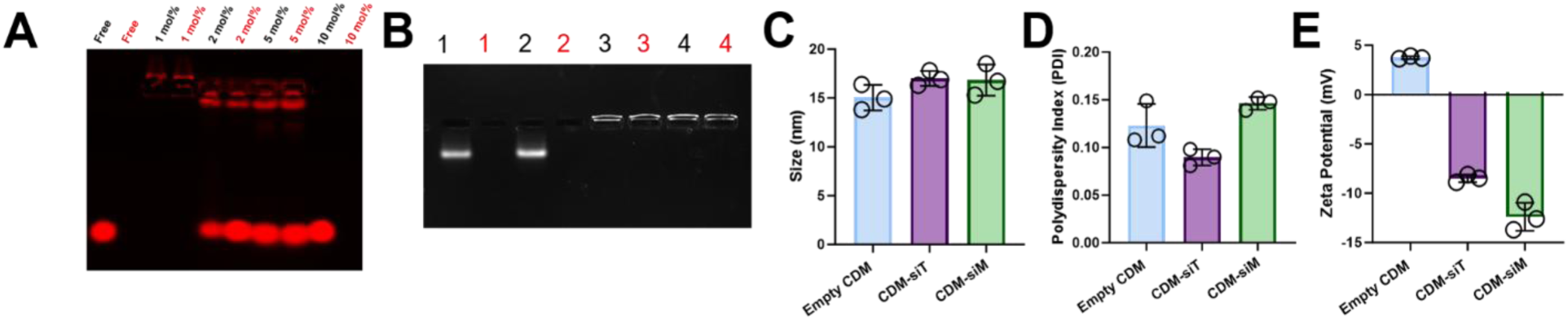
Characterization of siRNA-loaded CD-targeted micelles. **(A)** Gel shift assay showing incorporation of siRNA into CDM. Lanes were loaded with 500 ng of 21-mer siRNA under the indicated conditions. CDM formulated at 1 mol% prevented siRNA migration, consistent with stable incorporation. Lanes labeled in red were treated with RNase A prior to electrophoresis. **(B)** Gel shift assay demonstrating incorporation of TMEM16a siRNA or MCP-1 siRNA into CDM. Lanes are as follows: (1) Free *Tmem16a* siRNA, (2) free *Mcp-1* siRNA, (3) CDM-siT, and (4) CDM-siM. Red text indicates RNase A treatment prior to electrophoresis. **(C)** DLS analysis of hydrodynamic diameter for CDM, CDM–siT, and CDM–siM formulations. **(D)** Polydispersity index (PDI) of micelle formulations. **(E)** Zeta potential measurements of CDM and siRNA-loaded micelles.(n ≥ 3) Data are presented as mean ± SD. Statistical significance was determined by one-way ANOVA with Tukey’s post hoc test. * p < 0.05, **** p < 0.0001, ns = not significant.

**Figure S4:**
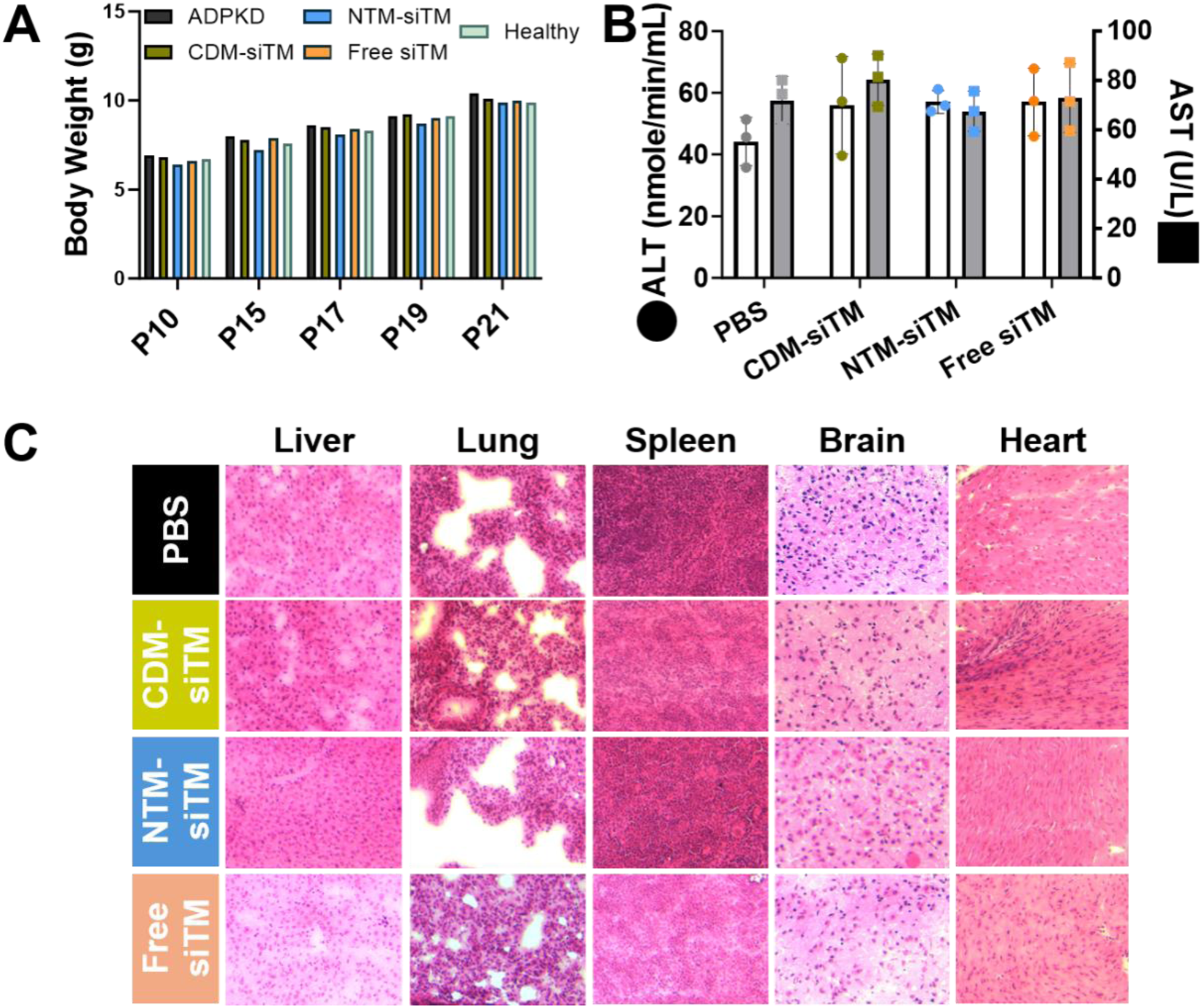
Liver enzymes and extrarenal organ morphology after repeated administration of CDM-siTM. **(A)** Body weight of mice over the course of treatment. **(B)** Serum analysis of liver function (AST and ALT) markers following administration of CDM-siTM at indicated doses compared to PBS and uninduced (healthy) controls, showing no significant changes across groups. **(C)** Representative H&E staining of major organs (liver, lung, spleen, brain, heart) after treatment. No overt histopathological abnormalities were observed across treatment groups. Data are presented as mean ± SD.

**Fig. S5:**
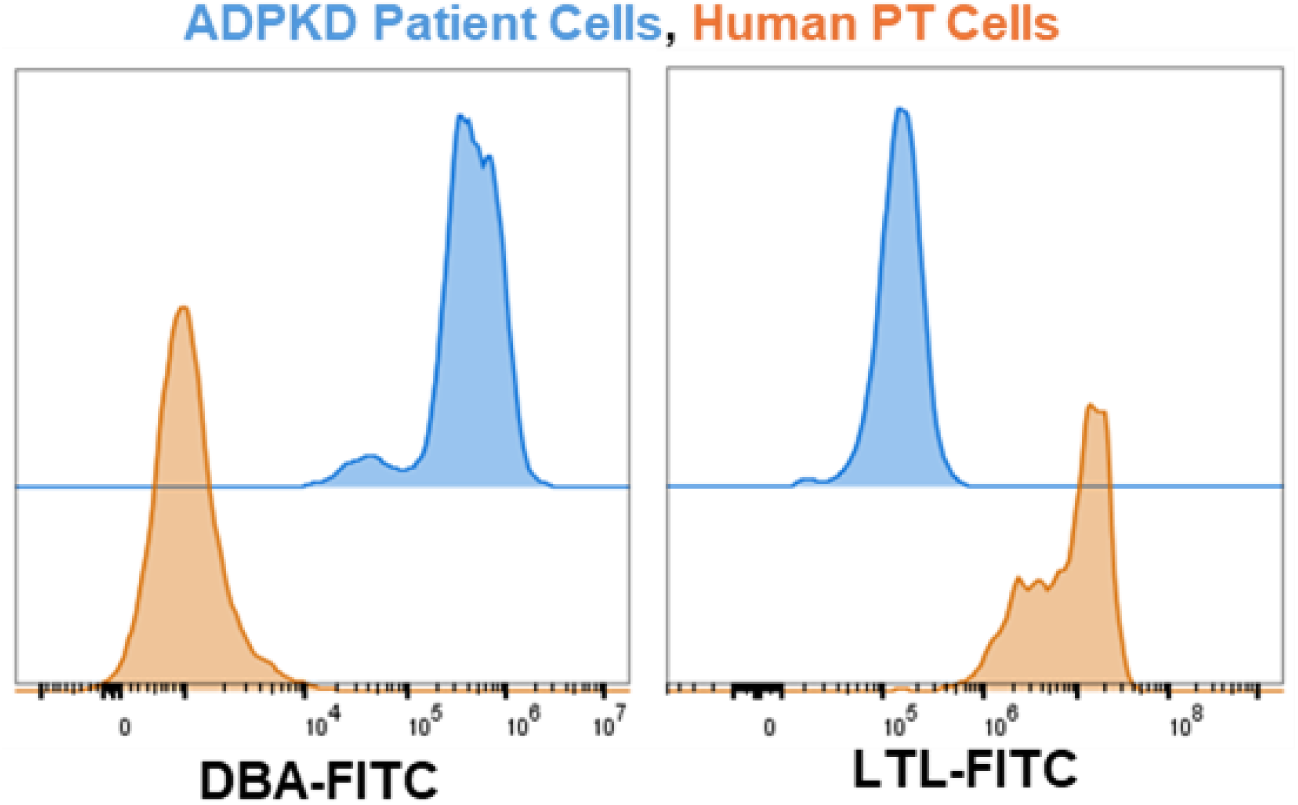
Characterization of ADPKD patient cells. Representative histograms of patient cells (blue) and human renal proximal tubular epithelial cells (human PT cells) stained with DBA-FITC or LTL-FITC. N = 3.

